# Phylogenomic subsampling and upsampling for efficient evolutionary analyses of big data

**DOI:** 10.64898/2026.06.21.733599

**Authors:** Sudhir Kumar, Koichiro Tamura, Sudip Sharma

## Abstract

Long runtimes, high memory demands, and reliance on high-performance computing impede phylogenomic analyses. We review a scalable phylogenomic subsampling with upsampling (PSU) framework, in which small subsamples of sites from a concatenated alignment are expanded by upsampling before inference, and the resulting analyses are then aggregated to obtain evolutionary estimates. PSU harnesses the fact that the computational cost of maximum likelihood analysis is strongly influenced by the number of distinct site patterns in the concatenated alignment, whereas statistical power depends primarily on the amount of evolutionary information represented by the total number of sites and substitutions. By reducing the former while restoring the latter through upsampling, PSU can approximate many full-data analyses at substantially lower computational cost. Analysis of simulated and empirical datasets shows that PSU can accurately estimate bootstrap support values, select the optimal substitution model, test evolutionary hypotheses, and infer branch lengths, divergence times, and associated uncertainty measures, while reducing runtime and memory requirements by orders of magnitude. PSU also provides distributions of inferred clade support across independent subsamples, enabling detection of conflicting phylogenetic signals that may remain hidden in conventional bootstrap analysis. Automated tuning of subsample size, the number of subsamples, and the number of upsampling replicates make PSU practical across diverse datasets. We suggest that PSU is a general strategy for scalable phylogenomic inference using a broad range of statistical methods. By enabling analyses of genome-scale alignments on commodity hardware, PSU broadens research access and reduces environmental and infrastructural costs of big-data phylogenomics.

## Introduction

Advances in sequencing technology and the rapid growth of sequence databases have made it increasingly easier to assemble large collections of sequences spanning thousands of loci across diverse organisms, individuals, and strains (**Fig. 1**). This expansion has transformed molecular phylogenetics into phylogenomics, which involves analyzing genome-scale alignments to infer evolutionary relationships, estimate divergence times, and investigate patterns of molecular evolution (Philippe et al. 2005; Kumar et al. 2012).

**Figure 1.**
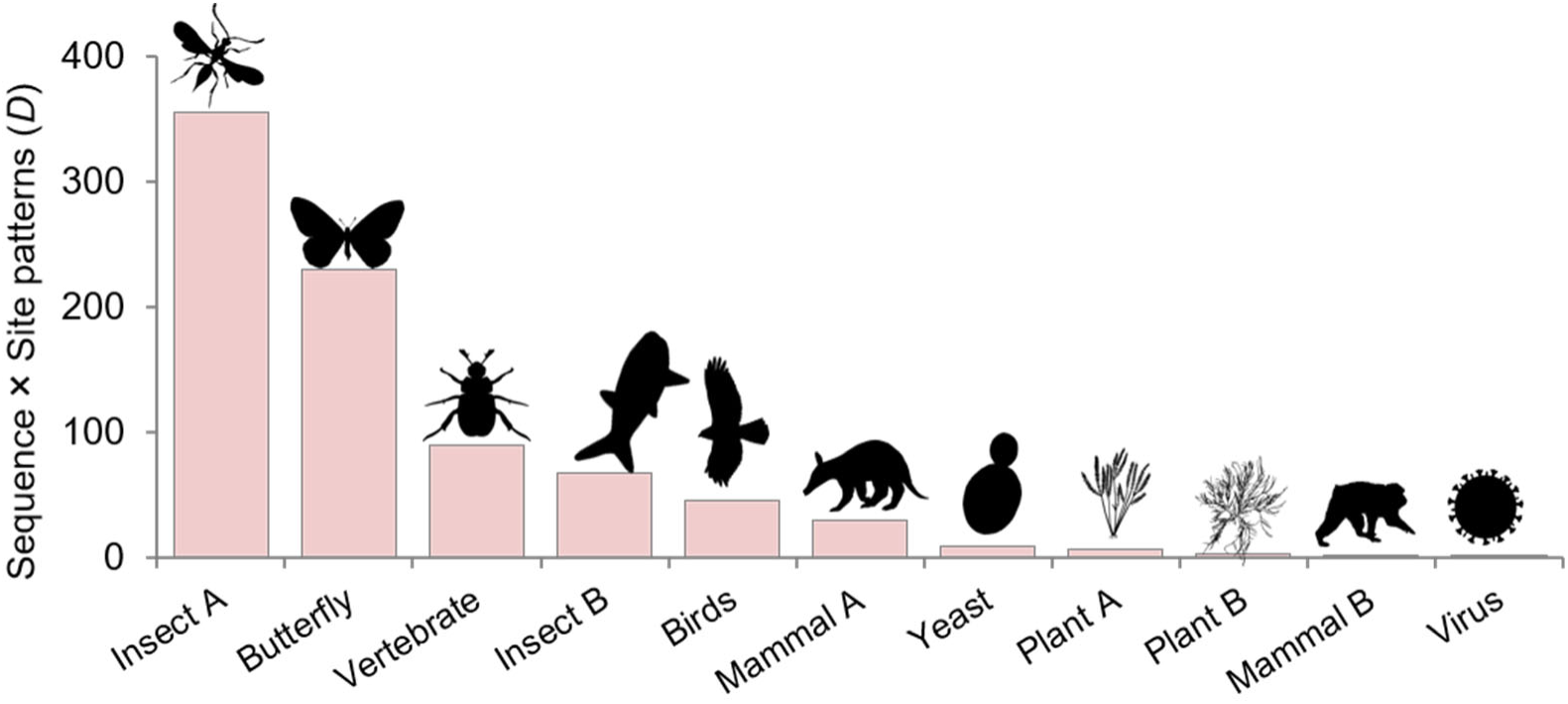
Phylogenomic datasets across the Tree of Life. Each bar represents a phylogenomic dataset, with bar height proportional to the data size (*D*), defined as the product of the number of distinct site patterns and the number of sequences in the concatenated alignment. The computational resource requirements of ML analysis depend on both the number of sequences and the number of unique site patterns in the alignment. Dataset references are in *Supplementary Table S1*.

The success of phylogenomics rests on a simple principle: longer alignments contain more evolutionary information and generally provide greater statistical power for estimating phylogenies and evolutionary parameters. However, the computational demands of analyzing these datasets increase dramatically with alignment size. This burden is particularly acute for computationally intensive methods such as maximum likelihood (ML; **Fig. 2**) and other sophisticated approaches. For example, selecting the best-fit nucleotide substitution model for an alignment of 394 kilobase pairs from 200 bird species (hereafter, the 200×394 dataset) required more than 10 days of computation (258 hours) and 14 GB of memory (Prum et al. 2015; Sharma and Kumar 2022). Similar computational demands are experienced when ML and other sophisticated methods are used to infer phylogenies, evaluate bootstrap support, and estimate divergence times (Sharma and Kumar 2021; Kumar 2022; Sharma and Kumar 2022). As phylogenomic datasets continue to grow, many analyses become impractical on standard desktop computers and increasingly depend on specialized high-performance computing resources.

**Figure 2.**
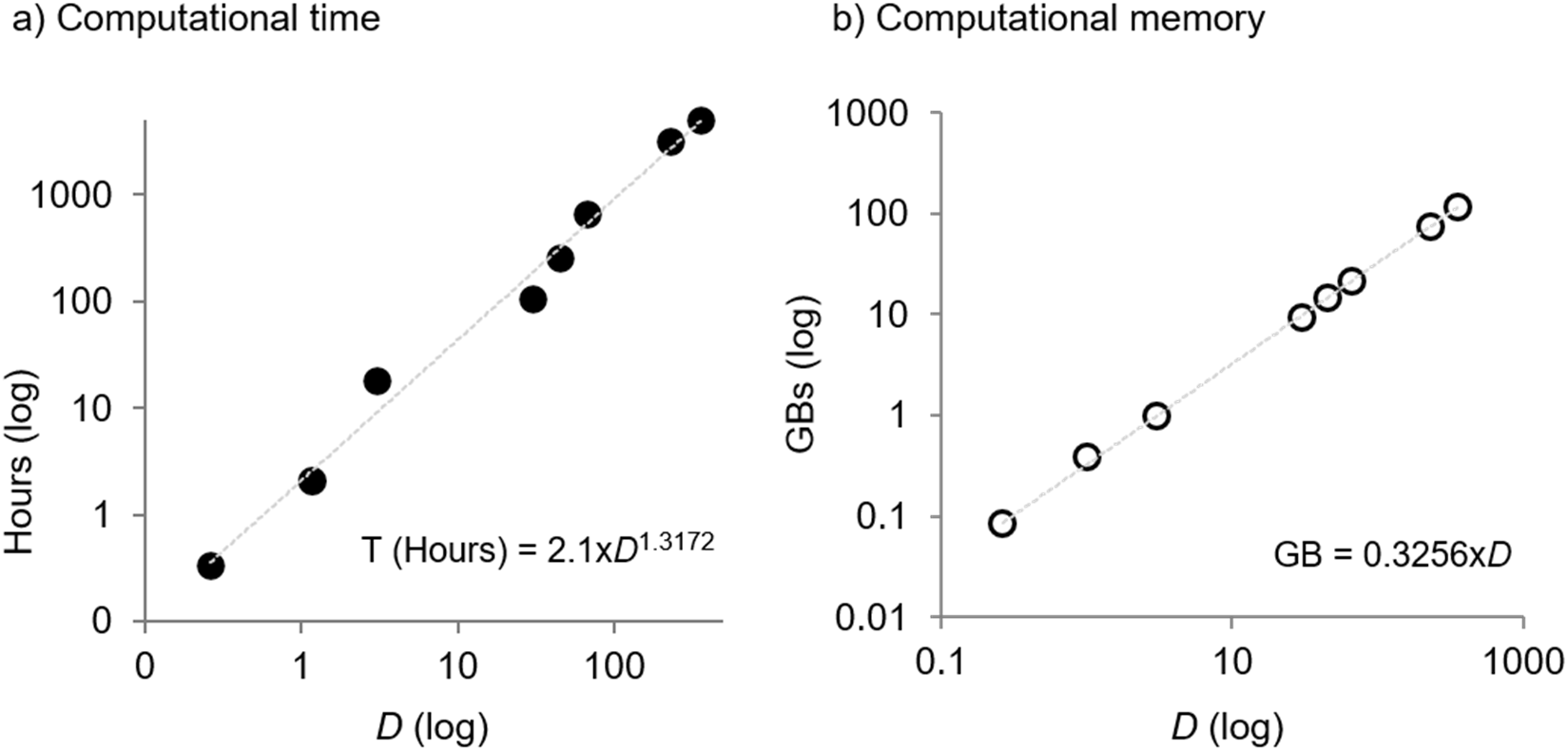
Increasing Computational demands. (**a**) Runtime and (**b**) memory requirements for ML analyses of eight empirical datasets, showing that computational resource requirements for model selection scale with the data size (*D*), which is the product of the number of sequences and the number of distinct site patterns. Source data are from Table 1 of Sharma and Kumar (2022).

**Table 1.** Performance of the PSU approach.

| Analysis type | PSU analysis |  |  |  |  | Full Data |  |
| --- | --- | --- | --- | --- | --- | --- | --- |
|  | Subsample size (max) | Savings |  | Time (seconds) | Memory (MBs) | Time (seconds) | Memory (MBs) |
|  |  | Time | Memory |  |  |  |  |
| Clade support | 2.1% | 79% | 96% | 994,142 | 144 | 4,734,008 | 3,773 |
| Model selection | 1.8% | 99% | 98% | 11,160 | 292 | 928,701 | 14,688 |
| Gamma parameter | 8.4% | 78% | 91% | 1,555 | 542 | 7,106 | 5,875 |
| Model substitution rates | 6.3% | 86% | 93% | 980 | 410 | 7,106 | 5,875 |
| Branch lengths (BLs) | 8.4% | 78% | 91% | 1,555 | 542 | 7,106 | 5,875 |
| Variance of BLs | 8.4% | 50% | 86% | 68,258 | 542 | 135,559 | 3,771 |
| Molecular Clock Test | 4.2% | 85% | 44% | 2,651 | 1,358 | 17,650 | 2,427 |

**Note.** The avian phylogenomic dataset comprised 259 nuclear loci from 200 species (Prum et al. 2015). The concatenated multiple sequence alignment contains 394,686 sites and 336,490 unique site patterns. All analyses were conducted using IQ-TREE 2 (Minh et al. 2020) with and without PSU.

These computational demands hinder discovery and reduce scientific rigor. Long runtimes discourage robustness checks across alternative models, assumptions, and data treatments, while reliance on specialized high-performance computing infrastructure limits accessibility for many researchers. Reproducibility also suffers when independent reanalyses require days of computation or more memory than is available on standard desktops. Furthermore, repeated large-scale analyses carry environmental costs because computational time translates into energy consumption and carbon emissions (Kumar 2022). Computational innovations are needed to make evolutionary analysis scalable for big-data phylogenomics.

A key observation that has motivated recent advances in scalable phylogenomics is that distinct properties of a concatenated phylogenomic alignment govern computational burden and inferential power. Computational cost depends strongly on the number of distinct site patterns across sequences analyzed, whereas statistical power depends primarily on the amount of evolutionary information represented by the total number of sites and substitutions. This distinction creates an opportunity: if one can preserve much of the statistical power contained in a large alignment while substantially reducing the number of site patterns analyzed, it may be possible to obtain accurate evolutionary inferences at a fraction of the computational cost.

The analysis of phylogenomic subsamples with upsampling (PSU) is an emerging framework that exploits this separation. In PSU, many random subsamples of alignment sites are drawn from the full concatenated alignment without regard to gene boundaries, partitions, or prior biological assumptions. Each subsample is then expanded via upsampling before statistical inference. This upsampling restores the full-alignment scale in terms of the total number of sites and substitutions represented, while retaining only a small fraction of the distinct site patterns found in the original dataset. Results from multiple subsamples are then aggregated to produce stable estimates. Because each analysis evaluates only a small subset of site patterns, computational requirements can be dramatically reduced without sacrificing statistical power.

Traditional phylogenomic subsampling approaches generally analyze only the selected subset of sites. Because these reduced datasets contain fewer sites and substitutions than the full alignment, they suffer a loss of statistical power and require additional assumptions to relate their estimates and variances to those obtained from the full dataset (Sharma and Kumar 2021; Sharma and Kumar 2022). So, instead of speeding up bootstrap analysis of long phylogenomic alignments, phylogenomic subsamples are frequently used to evaluate the stability of evolutionary inferences from concatenated datasets and to explore heterogeneity in phylogenetic signal across partitions and loci (Seo 2008; Faircloth et al. 2012; Song et al. 2012; Edwards 2016; Mongiardino Koch 2021; Lozano-Fernandez 2022).

It is worth noting that the PSU approach complements scalable bootstrap approaches designed to improve the efficiency of bootstrap analyses, such as rapid bootstrap procedures and approximate likelihood calculations (Stamatakis et al. 2008; Price et al. 2010a; Minh et al. 2013; Hoang et al. 2018). Those methods accelerate tree search and use approximate likelihood calculations to estimate bootstrap support, but they do not analyze subsamples of sites in a manner similar to PSU. Later, we discuss how those complementary approaches can be combined to achieve computational gains for phylogenomic datasets with many sites and many taxa.

While PSU builds on the statistical foundation of the bag-of-little-bootstraps framework (Kleiner et al. 2014), it extends it to the distinct needs of phylogenomics, such as selecting the substitution model and detecting conflicting phylogenetic signals (Sharma and Kumar 2021; Sharma and Kumar 2025). Here, we demonstrate that PSU can be applied to test evolutionary hypotheses and to estimate branch lengths, divergence times, and their variances. These developments establish that PSU is now more than a bootstrap acceleration technique. However, like many widely used strategies in phylogenetics, PSU should be viewed as a computational approximation rather than an exact formulation of full-data inference. This places it alongside heuristic tree-search methods, approximate likelihood calculations, and numerical optimization procedures that routinely trade computational efficiency for practical scalability, e.g., (Strimmer and Von Haeseler 1996; Price et al. 2010b; Stamatakis 2014; Minh et al. 2020; Azouri et al. 2021; Kumar et al. 2023; Kumar et al. 2024). The value of the PSU approach, therefore, rests on empirical performance, convergence behavior, and practical utility across realistic datasets. In the sections that follow, we review the logic, applications, performance, and limitations of PSU as a unifying framework for scalable phylogenomic inference.

### Estimating bootstrap support using PSU

Estimating confidence in inferred evolutionary relationships was the first major application of the PSU framework. Bootstrap support value estimation is particularly attractive because it requires repeated analyses of datasets nearly as large as the original alignment, thereby increasing computational demands rapidly with dataset size. PSU addresses this challenge by replacing full-alignment bootstrap analyses with analyses of many small site subsamples that are subsequently upsampled and aggregated.

#### Felsenstein’s bootstrap approach

Felsenstein (1985) adapted Efron’s (1979) bootstrap method to assess confidence in clades derived from molecular phylogenetic analysis. In Felsenstein’s bootstrap, *R* replicate alignments are generated by resampling sites with replacement from the full dataset of *N* sites. Each replicate dataset also has *N* sites. A phylogeny is inferred from each replicate dataset, and the proportion of replicates that reconstruct a particular clade is used as its full-alignment bootstrap support (*FBS*) (**Fig. 3a**). This analysis evaluates the statistical stability of inferred clades under sampling variation introduced by resampling sites from the full dataset. Felsenstein’s bootstrap procedure is computationally expensive because hundreds of phylogenetic analyses must be conducted on bootstrap replicate datasets, each containing ∼63.2% of the original sites. This is because the probability that a site is not chosen in a given draw is (1 - 1/*N*), and thus the probability that it is never chosen in all *N* draws is (1 - 1/*N*)*^N^*. Therefore, the probability that the site appears at least once is 1 - (1 - 1/*N*)*^N^*, which approaches 1 - *e*^-1^ ≈ 0.632 as *N* becomes large. Thus, each bootstrap replicate contains about 63.2% of the original sites and, thus, site patterns.

**Figure 3.**
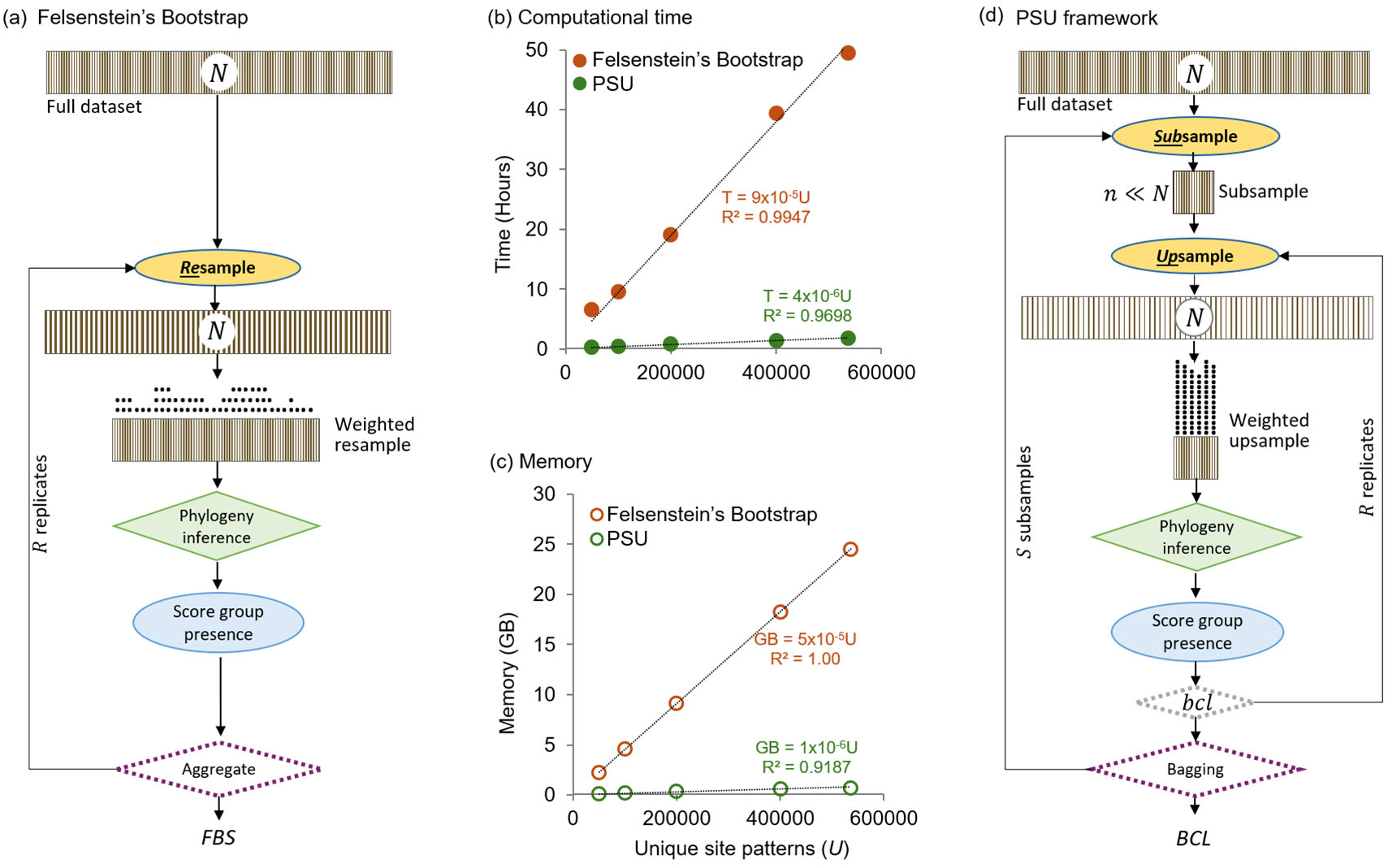
Flowcharts and computational demands of Bootstrap analysis. (**a**) In Felsenstein’s bootstrap, replicate datasets are generated by resampling sites from the full dataset (*N* sites and *U* distinct site patterns). Each bootstrap replicate dataset is subsequently compressed into a set of unique site patterns (*u*), with each site pattern assigned a weight equal to its frequency (stacks of dots) in the resampled dataset. This compressed representation is called a weighted resample. Full-alignment bootstrap support (*FBS*) for a clade is the proportion of replicate phylogenies containing the clade of interest. (**b**) runtime and (**c**) memory usage for inferring ML phylogeny for one bootstrap replicate for increasingly longer phylogenomic datasets. (**d**) In the PSU framework, small subsamples of randomly selected *n* sites (*n* << *N*) are upsampled to full length (*N* sites). The resulting upsampled datasets are then compressed into unique site patterns, each assigned a weight equal to its frequency in the upsampled dataset, thereby generating weighted upsamples for ML analysis. Weighted upsampled PSU datasets contain a small number of site patterns, each occurring at a much higher frequency than in *FBS* analysis. Clade support is estimated for each subsample (*bcl*), then aggregated across *S* subsamples to obtain the median *bcl* value (*BCL*), which is used to estimate *FBS*. Panels **b** and **c** show the time and memory required to analyze a PSU replicate dataset. Results are from the analysis of simulated datasets (Tamura et al. 2012) containing 446 sequences and alignments ranging from 50,000 to 536,534 bases, as reported in Fig. 1b of Sharma and Kumar (2021).

Since computational time scales linearly with the number of unique site patterns in the dataset (**Fig. 3b**), Felsenstein’s bootstrap analysis with just 100 replicates is expected to take more than 63 times as long as a single ML phylogeny inferred from the full dataset. These requirements escalate with more replicates and can become onerous for very long phylogenomic alignments (**Fig. 3b**). Memory requirements per replicate also increase linearly with the number of unique site patterns, which can far exceed the memory available on standard desktops (**Fig. 3c**). These demands motivated the need for alternative strategies that can generate accurate estimates of bootstrap support while greatly reducing computational costs.

#### PSU-based bootstrap support estimation

Sharma and Kumar (2021) adapted and advanced the bag-of-little-bootstraps approach of Kleiner et al. (2014) for phylogenomic analysis (**Fig. 3d**). Each PSU analysis begins by selecting a random subset of sites (*n* << N) from the full alignment. Bootstrap replicate datasets are generated by resampling *N* sites with replacement from that subsample. The number of sites in each upsampled replicate alignment, therefore, matches the full dataset length (*N*) while containing no more than *n* unique site patterns present in the subsample. This upsampling step is central to the PSU framework, which restores the statistical scale of the analysis without the computational burden of the full alignment. As shown in **Figure 4**, the number of substitutions per upsampled PSU replicate is comparable to that in conventional bootstrap replicates of the full dataset. Thus, PSU replicates trade total site diversity for computational efficiency while preserving much of the statistical scale needed for confidence estimation.

**Figure 4.**
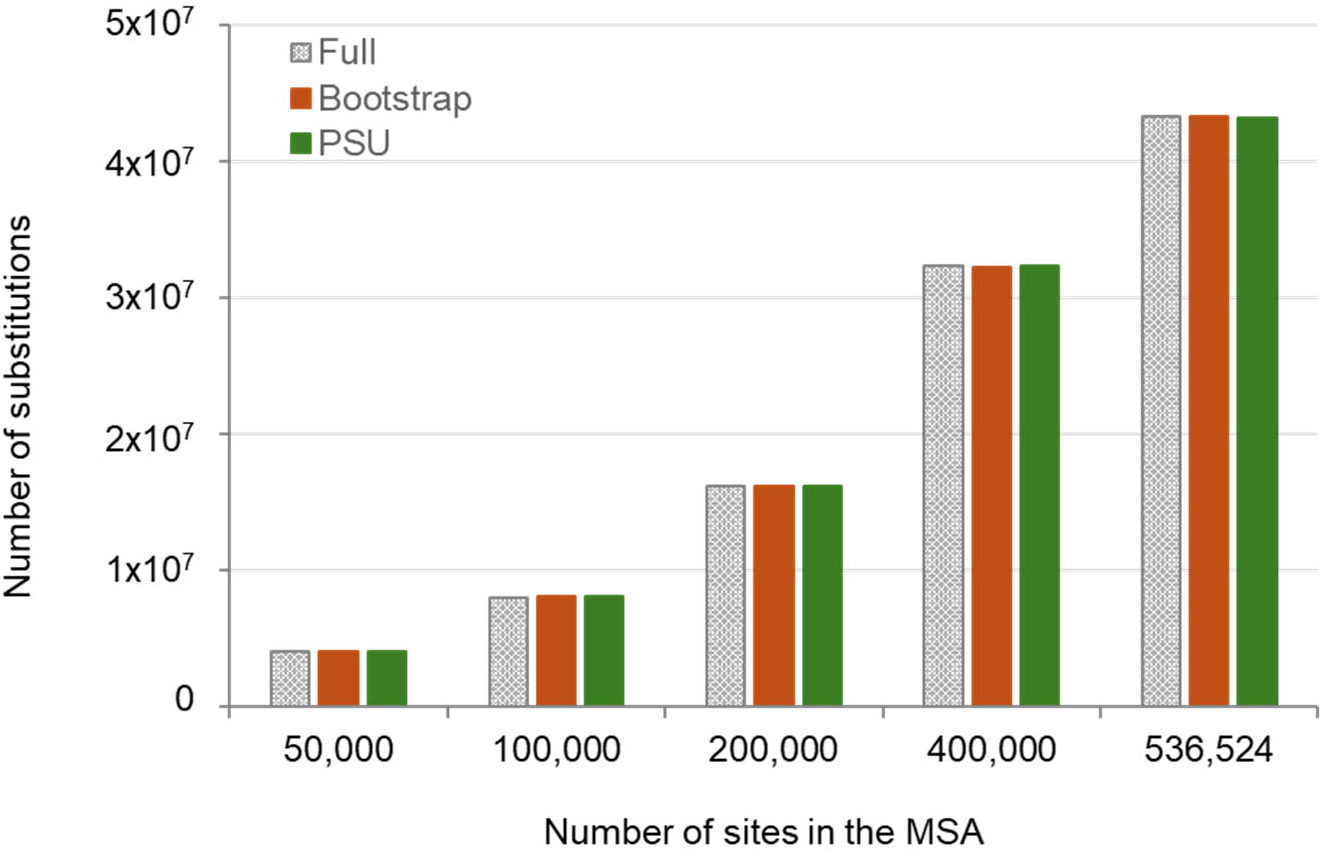
Number of substitutions in PSU datasets. The number of substitutions in PSU replicate datasets (green) as compared to those in the standard bootstrap replicate datasets (red) and the full dataset (grey). Results are from the analysis of five simulated datasets, containing 446 species and MSAs with 50,000 to 536,534 sites, as reported by Sharma and Kumar (2021). The number of substitutions is the sum of ML estimates of branch lengths multiplied by the number of sites.

In the PSU analysis, multiple upsampled datasets are generated from each subsample. A phylogeny is inferred for each PSU replicate, and the proportion of PSU trees containing a specific clade yields the subsample bootstrap confidence level (*bcl*). Because each subsample includes only a fraction of the sites in the full dataset, multiple subsamples must be analyzed to consider the diversity of evolutionary information in the full dataset, which is also needed to obtain a stable estimate of full-data bootstrap support. Thus, bcl values across subsamples are aggregated to estimate *FBS* for a given clade.

The way subsample *bcl* values are aggregated is important. Sharma and Kumar (2021) found that the mean *bcl*, although strongly correlated with *FBS*, tended to underestimate high *FBS* values and overestimate low *FBS* values. The median of the subsample *bcl* values, denoted *BCL*, provided a better approximation of *FBS*. This difference between mean and median *bcl* reflects the indicator nature of clade support. Each replicate tree either contains a clade or does not, unlike the continuous estimators emphasized in the original bag-of-little-bootstraps framework of Kleiner et al. (2014). The *Appendix* provides a simple illustration of why median outperforms mean aggregation for such indicator-style measurement of confidence levels.

*BCL* estimation requires only a fraction of the time and memory needed for full-data bootstrap analysis. For the 200×394 dataset, PSU reduced computation time by 79% and memory use by 95% (**Table 1**), with similar gains reported for other empirical datasets (Sharma and Kumar 2021). In many cases, subsampling fewer than 5% of sites was sufficient, and approximately 10 subsamples, each with about 10 upsampling replicates, produced stable results. These reduced memory demands also make PSU analyses easier to parallelize across cores, further decreasing wall-clock time. PSU is, therefore, especially useful when full-data bootstrap analysis would exceed desktop memory limits or require days of computation.

#### Distributions of clade support from PSU

Unlike full-data bootstrapping, which reports a single support value for each clade (*FBS*), PSU produces a distribution of support values across subsamples for the selected subsample size (*n*). Some clades show consistently high support across subsamples, indicating strong genome-wide concordance (**Fig. 5a**). Others show multimodal distributions, with some subsamples supporting the inferred clade and others contradicting it (**Fig. 5b**). Such heterogeneity can reflect alignment or orthology errors, model misspecification, hidden biases, incomplete lineage sorting, or other sources of phylogenetic discordance (Lanfear and Hahn 2024; Sharma and Kumar 2024; Sharma and Kumar 2025). PSU detects this heterogeneity directly from random site subsamples rather than from predefined genes or partitions.

**Figure 5.**
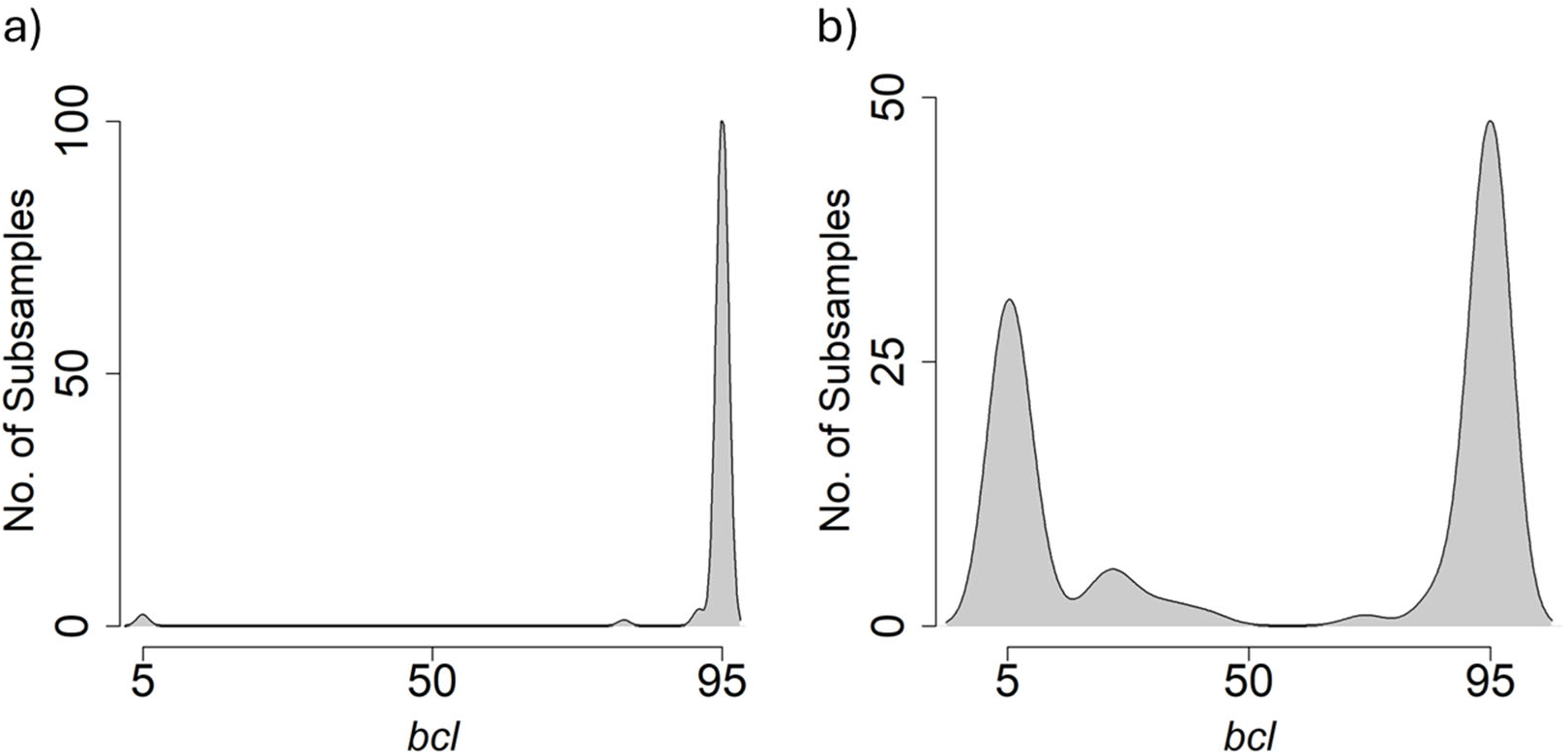
Distribution of subsample bootstrap support values (*bcl*) for two rodent clades. (**a**) Consistently high support across subsamples for a clade. (**b**) A distribution indicating conflicting phylogenetic signals across subsamples for the inferred clade. Results are for clades R1 and R1’, respectively, in the rodent phylogeny presented in Figure 3 of Sharma and Kumar (2025).

The distribution of *bcl* values can be used to estimate Net Bootstrap Support, *NBS*, a measure designed to reduce overconfidence in concatenated phylogenomic analyses (Sharma and Kumar 2025). *NBS* is calculated as the mean of the subsample *bcl* distribution, whereas *BCL* uses the median *bcl* to approximate full alignment bootstrap support (*BCL* ∼ *FBS*). This distinction is important. The median is useful for approximating *FBS*, but it downplays conflict among subsamples. The mean incorporates the entire distribution, including subsamples that weakly support or contradict a clade, and, therefore, provides a theoretically supported measure that is more sensitive to phylogenetic conflict (Sharma and Kumar 2022).

Analyses of some simulated datasets suggest that *NBS* can perform similarly to multispecies coalescent (MSC) methods across phylogenomic datasets generated under low, moderate, and high levels of ILS (**Fig. 6a**) (Sharma and Kumar 2025). The performance of *NBS* was also evaluated under conditions of gene tree estimation error (GTEE), which is another major source of gene tree discordance (Degnan and Rosenberg 2009; Simmons and Gatesy 2015; Mirarab et al. 2016; Shen et al. 2021). GTEE arises from inaccurate estimation of gene trees due to short gene alignments or limited phylogenetically informative substitutions, which can influence both tree inference and clade-support estimation (Shen et al. 2021; Sharma and Kumar 2025). Interestingly, *NBS* could surpass MSC-based local posterior probabilities when gene tree estimation error (GTEE) was high (**Fig. 6b**). This is because MSC methods are impacted by poor resolution in the input gene trees (Simmons and Gatesy 2015; Shen et al. 2021; Sharma and Kumar 2025), whereas *NBS* estimation does not involve gene tree estimation.

**Figure 6.**
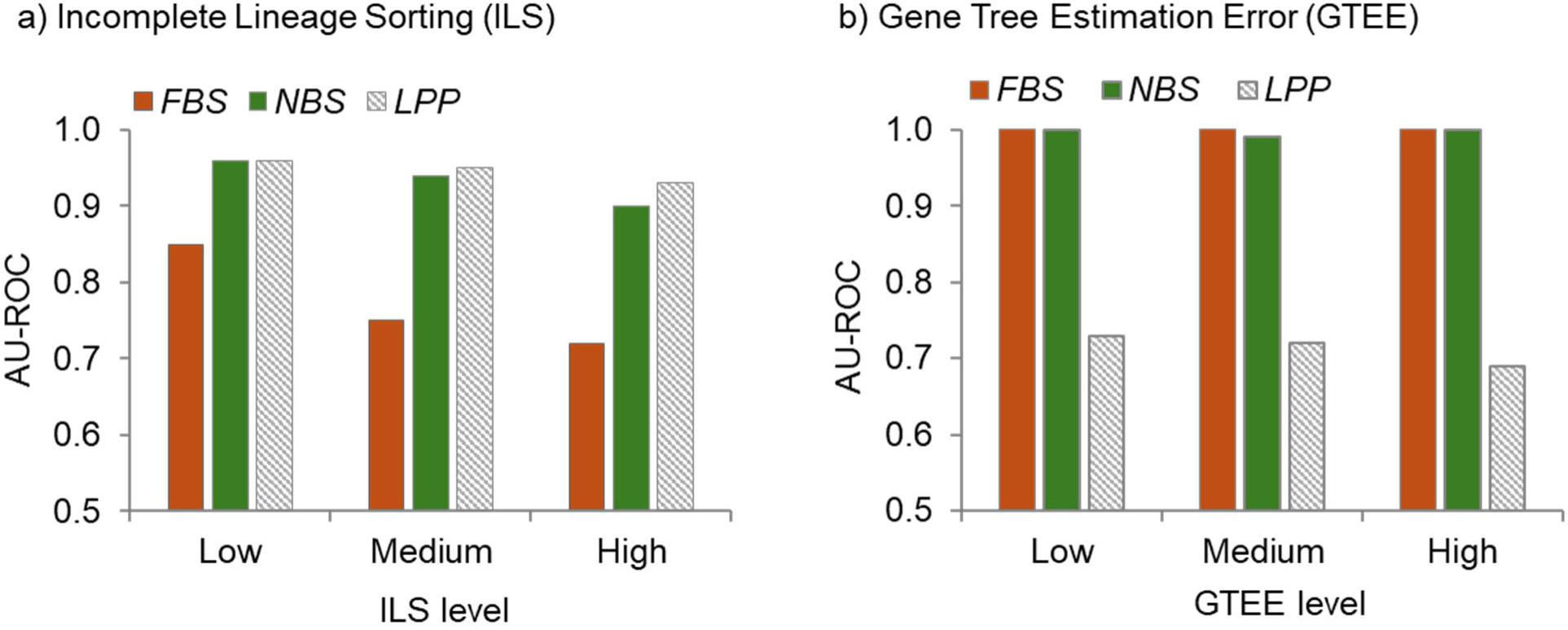
Accuracy of phylogeny support measures under heterogeneous conditions. Performance of support metrics in distinguishing correct and incorrect clades under varying levels of (**a**) incomplete lineage sorting (ILS) and (**b**) gene tree estimation error (GTEE), redrawn using results in Figures 7 and 8 of Sharma and Kumar (2025). The performance was evaluated using the area under the receiver operating characteristic curve (AU-ROC), which quantifies the ability of a support metric to correctly discriminate between correct and incorrect clades at different bootstrap value thresholds. AU-ROC is 0.5 when the metric’s performance is no better than random designation. Simulated datasets were obtained from Mirarab et al. (2014) and simulated with low, medium, and high levels of ILS using a species tree of 37 mammalian species within a multispecies coalescent framework. Ten datasets were analyzed from each ILS category, each comprising 100 genes (1,600 sites per gene). Simulated datasets with GTEE were obtained from Shen et al. (2021), in which branch lengths were reduced to introduce low, medium, and high GTEE levels, with 1,000 gene sequence alignments for each category. Net Bootstrap Support (*NBS*), derived from the mean of subsample support values, incorporates both supporting and conflicting signals and achieves performance comparable to or exceeding local posterior probability (*LPP*) produced by MSC-based approaches under challenging conditions.

In addition to ILS and GTEE, one or a few loci with unusual phylogenetic histories, alignment issues, and hidden biases can drive bootstrap support for incorrect or contentious clades in concatenated analyses of hundreds of loci (Brown and Thomson 2016; Shen et al. 2017; Sharma and Kumar 2024; Sharma and Kumar 2025). In the PSU analysis, some subsamples will exclude such disruptive loci and sites, thereby reducing support for spurious clades driven by those sites (Sharma and Kumar 2025). In these cases, *NBS* is expected to be lower than *FBS*, as observed in the concatenated analysis of multiple empirical datasets (Sharma and Kumar 2025). Thus, *NBS* can serve as a practical way to reduce overconfidence while simultaneously revealing heterogeneity in phylogenetic signals across the alignment.

#### Phylogenomics without data partitions

Conventional approaches for addressing phylogenetic heterogeneity divide an alignment by gene, codon position, genomic region, or other biologically defined categories. These partitions may then be modeled separately in concatenated analyses or used to reconstruct gene trees, which are summarized using consensus or multispecies coalescent methods (Bull et al. 1993; Chippindale and Wiens 1994; Brandley et al. 2005; Gadagkar et al. 2005; Lanfear et al. 2017). Such strategies are powerful and widely used, but they also require decisions about how to partition the data, and different partitioning schemes can lead to different conclusions. In addition, fitting separate models to many partitions or estimating many gene trees can be computationally demanding, and short partitions may suffer from large GTEEs (Simmons and Gatesy 2015; Shen et al. 2021; Sharma and Kumar 2025). PSU offers a complementary strategy. Rather than requiring predefined partitions, it uses random subsamples of sites to assess whether support for a clade is consistent across the alignment. But this should not be taken to replace model-based approaches that explicitly assess gene-tree variation.

#### Relationships among FBS, BCL, and NBS

*FBS*, *BCL*, and *NBS* quantify related but distinct properties of phylogenetic support. *FBS* measures the frequency with which a clade appears in bootstrap replicates generated from the full alignment. *BCL* is designed to approximate this quantity by the median of the support values obtained from many independent PSU subsamples (Sharma and Kumar 2021). *NBS*, in contrast, measures mean support across subsamples and, therefore, reflects the consistency of phylogenetic signals throughout the alignment for a given subsample size.

Even when *BCL* (median, black circles) stabilizes and becomes the same as *FBS* at a given subsample size, *NBS* (mean, green circles) can be substantially lower than *BCL* (**Fig. 7a**). Doubling the number of sites in the subsample will increase *NBS*, making it closer to *BCL* that remains unchanged because individual subsamples become increasingly representative of the full concatenated alignment and disagreement among subsamples decreases (**Fig. 7a**). Consequently, the number of inferred clades supported with high *NBS* also keeps increasing, becoming similar to *BCL* as the number of sites sampled are increased (**Fig. 7b**). So, *NBS* is best estimated for minimum subsample size at which *BCL* stabilizes during adaptive PSU analysis (Sharma and Kumar 2021), which is also described below. We therefore suggest reporting NBS for the automatically selected subsample size (see below) alongside *BCL* in analyses of concatenated phylogenomic datasets.

**Figure 7.**
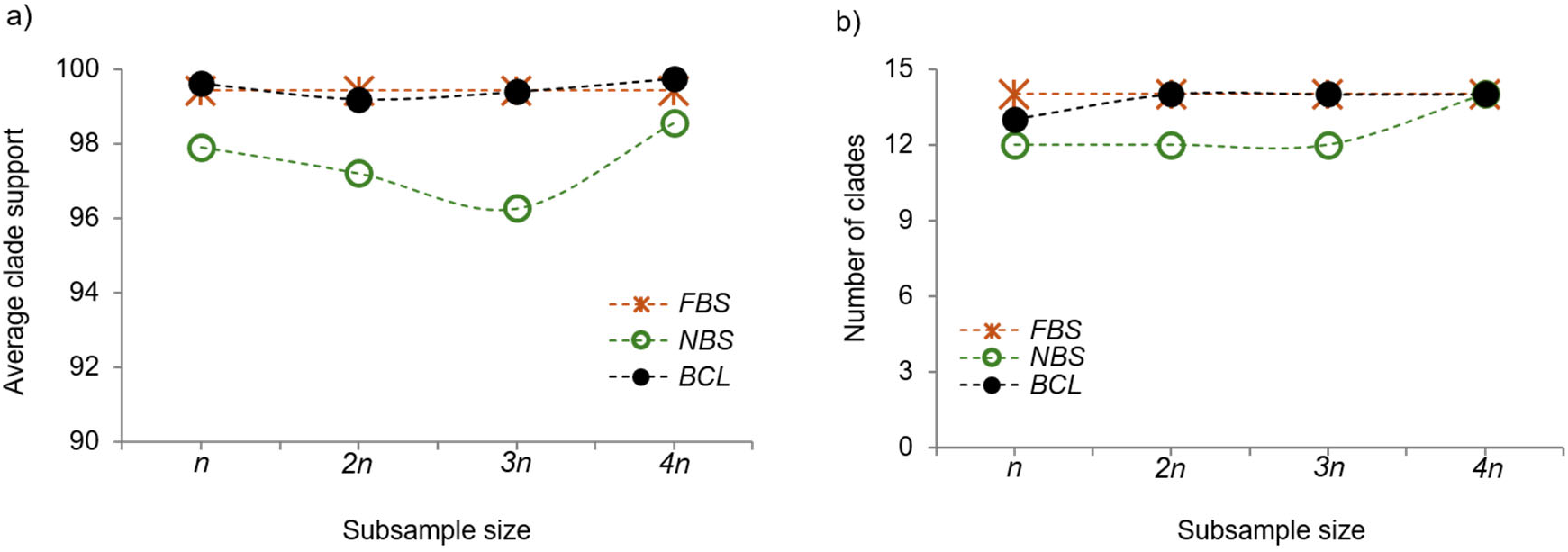
Trends of average *BCL* and *NBS* with increasing subsample size. (**a**) Average *BCL* (black) and *NBS* (green) using increasingly larger subsamples (*n* = 33,495) for clades with *FBS* ≥ 90% (orange). (**b**) Number of clades with *FBS* ≥ 90% recovered by *FBS*, *BCL,* and *NBS* using different sample sizes. Results were generated from a phylogenomic alignment of 20 species and all 394,684 sites from the 200x396kbp dataset. All analyses were conducted using IQTREE 2 with and without PSU.

#### Autotuning PSU analysis

PSU analysis requires choices for three tuning parameters: the number of sites per subsample (*n*), the number of independent subsamples (*S*), and the number of upsampling replicates per subsample (*R*). These choices affect both accuracy and computational efficiency, so adaptive protocols have been developed to select them objectively (Sharma and Kumar 2021; Sharma and Kumar 2025). The general strategy is to begin with an empirically guided subsample size and small numbers of initial subsamples and upsamples (Sharma and Kumar 2021). It was found that initial subsample size selection depends on the number of unique site patterns in the dataset, and this selection can be further refined by accounting for the number of taxa (Kumar *et al*., in preparation).

Support values are then estimated and monitored across iterations. Additional subsamples are added until average support values (*BCL* or *NBS*) for clades in different support ranges stabilize within predefined tolerances. If convergence is not achieved, the subsample size is increased (2*n*, 3*n*, …), and the process is repeated. In this case, *BCL* is used for convergence assessment across subsample sizes, rather than *NBS*, because *NBS* will generally increase with longer subsample sizes (**Fig. 7a-b**). Once stability criteria are met, *BCL* and *NBS* are estimated from the resulting collection of PSU replicate phylogenies. This adaptive approach reduces the need for *ad hoc* parameter tuning and makes PSU analyses more reproducible. It is also practically important because optimal choices depend on dataset-specific properties such as alignment length, divergence, missing data, and phylogenetic complexity.

### Evolutionary Hypothesis Testing using PSU

The applications discussed thus far focused on bootstrap support estimation for phylogenetic relationships. However, the PSU framework is not limited to bootstrap support. Many core tasks in evolutionary biology involve selecting among competing models, testing evolutionary hypotheses, and comparing alternative explanations for observed sequence variation. These analyses often require repeated likelihood calculations on large alignments and can become major computational bottlenecks in phylogenomics. PSU provides a general strategy for these problems: compute the relevant likelihood-based statistic on upsampled subsamples and aggregate results across subsamples until stable conclusions are obtained. In this way, PSU extends naturally from confidence estimation to a broader class of evolutionary inference problems.

#### Selecting the optimal substitution model

Choosing nucleotide or amino acid substitution models is a standard step in likelihood-based phylogenetics, typically involving likelihood-ratio tests or information criteria (Posada and Crandall 1998; Posada 2008; Kalyaanamoorthy et al. 2017; Darriba et al. 2020; Sharma and Kumar 2022). Model selection is particularly attractive for PSU because the inferential goal is discrete, the identity of the preferred model, rather than estimation of a continuous parameter, allowing convergence to be assessed directly through agreement among successive analyses.

For long phylogenomic alignments, this step can become expensive because many candidate models must be optimized and compared. PSU reduces this cost by evaluating candidate models on upsampled subsamples rather than on the full alignment (Sharma and Kumar 2022). For the 200×394 dataset, PSU selected the same final model as in the full-alignment analysis, reducing runtime from 258 CPU hours to 1.5 CPU hours and memory use from 14 GB to 0.15 GB (**Table 1**). Across additional empirical datasets, PSU showed orders-of-magnitude reductions in memory use and substantially improved scaling with data size (**Fig. 8b** and **c**). For model selection, only one upsampled replicate is needed for each subsample size, since the goal is not to estimate bootstrap support. The subsample size is increased progressively until the same substitution model is selected in consecutive analyses (**Fig. 8a**) (Sharma and Kumar 2022). In empirical applications, convergence was usually achieved using less than 5% of the unique site patterns in the full dataset.

**Figure 8.**
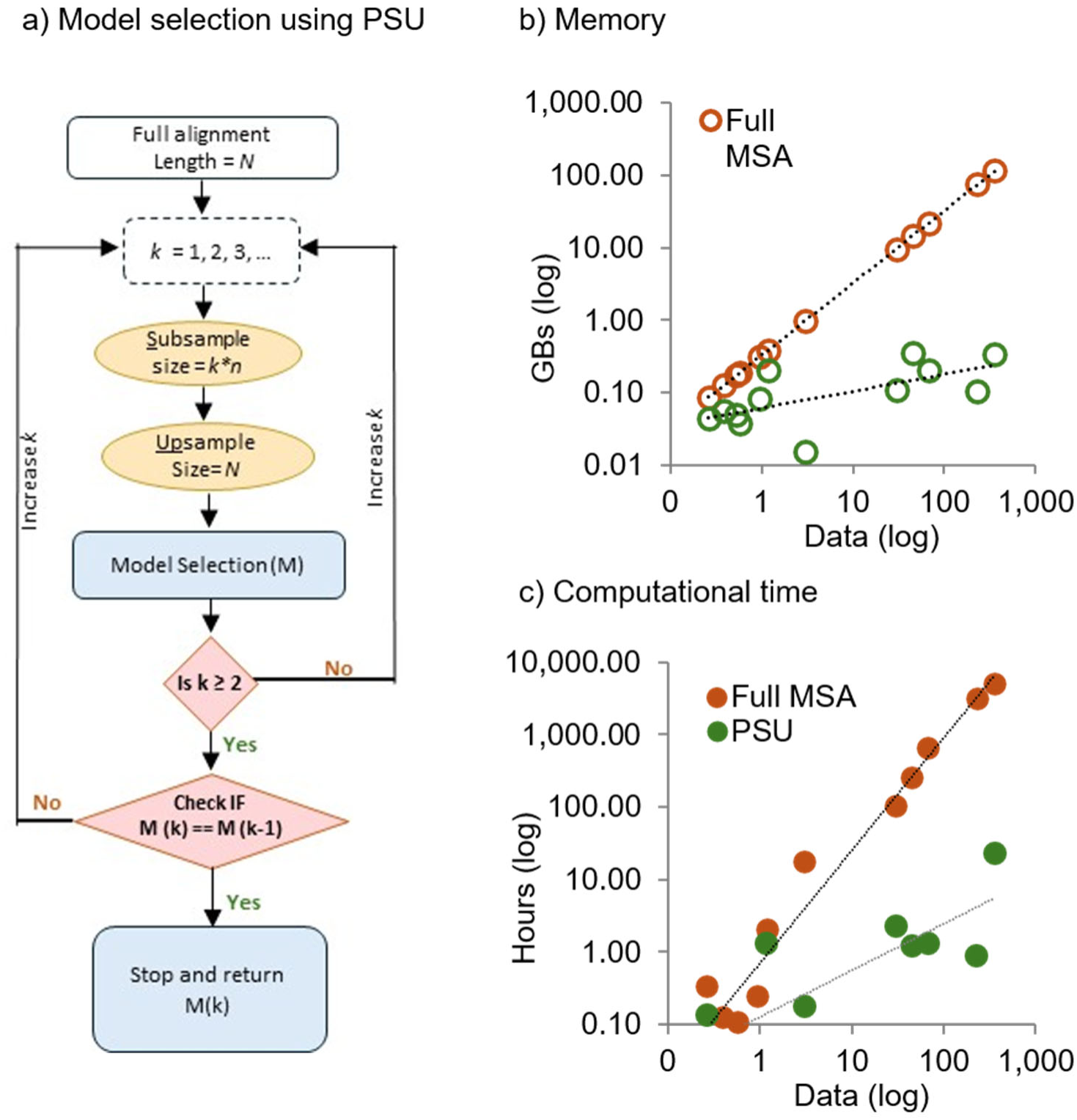
Scaling behavior of PSU versus full-data analysis for model selection. (**a**) A flowchart demonstrating the steps of the PSU framework for selecting the best-fit model of substitution. (**b**) Memory and (**c**) runtime requirements for substitution model selection as a function of data size (*D*, log scale), using data from Sharma and Kumar (2022). PSU exhibits sublinear scaling because the number of unique site patterns evaluated remains limited, even as the total number of sites and substitutions scales to full dataset size through upsampling. PSU time and memory needs are of order *O*(*D*^0.64^) and *O*(*D*^0.23^), respectively, compared to the full-data analysis, which are of order *O*(*D*^1.56^) and *O*(*D*^1.0^), respectively, for the dataset analyzed (see Figure 2).

#### Testing nested evolutionary hypotheses

More broadly, many evolutionary questions are formulated as comparisons between nested models that differ in one or more biological assumptions. Examples include molecular clock tests, local clock models, constraints on substitution processes, and models of selective pressure. Because these analyses ultimately depend on likelihood-ratio statistics, they are natural candidates for the PSU framework. For example, the molecular clock hypothesis can be tested by comparing the log-likelihood of a rooted phylogeny under clock and no-clock models and computing the usual test statistic, 2Δ*lnL*. In a PSU implementation, the test is first conducted on an upsampled subsample. The subsample size is then progressively increased until consecutive analyses yield the same statistical conclusion at a chosen significance threshold. The final test statistic and P-value are taken from the last converged analysis. Applied to the 200×394 dataset, this protocol converged using only 4.2% of the full dataset, and produced the same biological conclusion as the full-data analysis, rejecting the molecular clock with a highly significant *P*-value (**Table 1**).

Although illustrated here with the molecular clock test, the same strategy can be applied to other nested evolutionary hypotheses, including local-clock models and nested constraints on substitution processes or selection regimes, e.g., (Yoder and Yang 2000).

PSU therefore provides a scalable route to likelihood-based hypothesis testing when full-data calculations may be time-consuming or impractical.

#### Testing non-nested hypotheses

Non-nested comparisons constitute a particularly important class of phylogenomic problems because competing hypotheses often correspond to alternative species relationships or evolutionary scenarios that cannot be compared using standard likelihood-ratio tests. In such cases, inference depends on estimating the distribution of a test statistic rather than simply selecting the model with the highest likelihood. PSU can provide an efficient framework for estimating these distributions while avoiding repeated analysis of the full alignment.

For example, PSU can be used to test tree topologies (e.g., T1 and T2). In a typical bootstrap setup, the distribution of *Δln L* = *ln L*(*T*1) − *ln L*(*T*2) can be computed across bootstrap replicates under a fixed substitution model. The null hypothesis *Δln L* = 0 can be tested by constructing a confidence interval for the statistic using PSU, based on the estimated variance of *Δln L*. We conducted such an analysis for two contrasting rodent trees: one from a concatenated alignment (T1) and the other from an MSC analysis (T2). They differ in the placement of a single taxon (Roycroft et al. 2020; Shen et al. 2021; Sharma and Kumar 2025).

We estimated the variance of *Δln L* across multiple upsampling replicates for each subsample, then averaged these estimates across subsamples, following the logic of Kleiner et al. (2014). Additional subsamples were added until the variance estimate stabilized within a predefined tolerance. In this example, convergence was achieved with seven subsamples. The resulting 95% confidence interval for *Δln L* ranged from -78.6 to 231.3, including 0, indicating that the concatenation tree was not significantly better supported than the alternative topology. This result is consistent with the low *NBS* for the defining clade of T1 and with its low posterior probability in the MSC analysis (Roycroft et al. 2020; Shen et al. 2021; Sharma and Kumar 2025).

These applications demonstrate that PSU is not restricted to confidence estimation. It can be used whenever the objective is to select among competing evolutionary models, test biological hypotheses, or estimate the uncertainty associated with statistical estimates. This flexibility suggests that PSU should be viewed as a general framework for scalable evolutionary inference rather than solely as an alternative approach to bootstrap analysis.

### Estimating Evolutionary Parameters using PSU

The applications discussed thus far involve confidence estimation, model selection, and hypothesis testing. A more stringent evaluation of the PSU framework is whether it can accurately estimate continuous evolutionary parameters. Evolutionary parameter estimates are numerical quantities whose precision depends on both sampling variance and model assumptions, so the successful application of PSU to branch-length and divergence-time inference provides an important demonstration that it can support a broad range of phylogenomic inference tasks.

#### Estimating substitution model parameters

Substitution-model parameters estimated during PSU model selection were reported to be nearly identical to those obtained from full-data analysis, including the gamma shape parameter for among-site rate variation (Sharma and Kumar 2022). This suggests that PSU can be used not only to select models but also to efficiently estimate their parameters. For example, PSU can estimate the gamma shape parameter (*α*) by first analyzing a subsample with *n*, determined using previous approaches (Sharma and Kumar 2022), and then progressively increasing the subsample size and monitoring changes in α across iterations. The process stops when consecutive estimates fall within a predefined tolerance, 1% by default.

Applied to the 200×394 dataset, this PSU protocol converged after the fourth iteration, using at most 8.38% of all sites, and required less memory and runtime than full maximum-likelihood estimation using the complete dataset. Across empirical datasets, PSU-based estimates of *α* (0.398) closely matched full-data estimates (0.394). The same strategy can be used for substitution-rate parameters. In this case, convergence is assessed by comparing rate estimates across consecutive iterations, for example, by requiring a correlation coefficient of at least 0.99. For the 200×394 dataset, convergence was achieved after the third iteration using only 6.3% of sites, and the resulting rate estimates closely matched those from the full alignment (**Fig. 9a**; slope = 1.00, correlation = 0.999).

**Figure 9.**
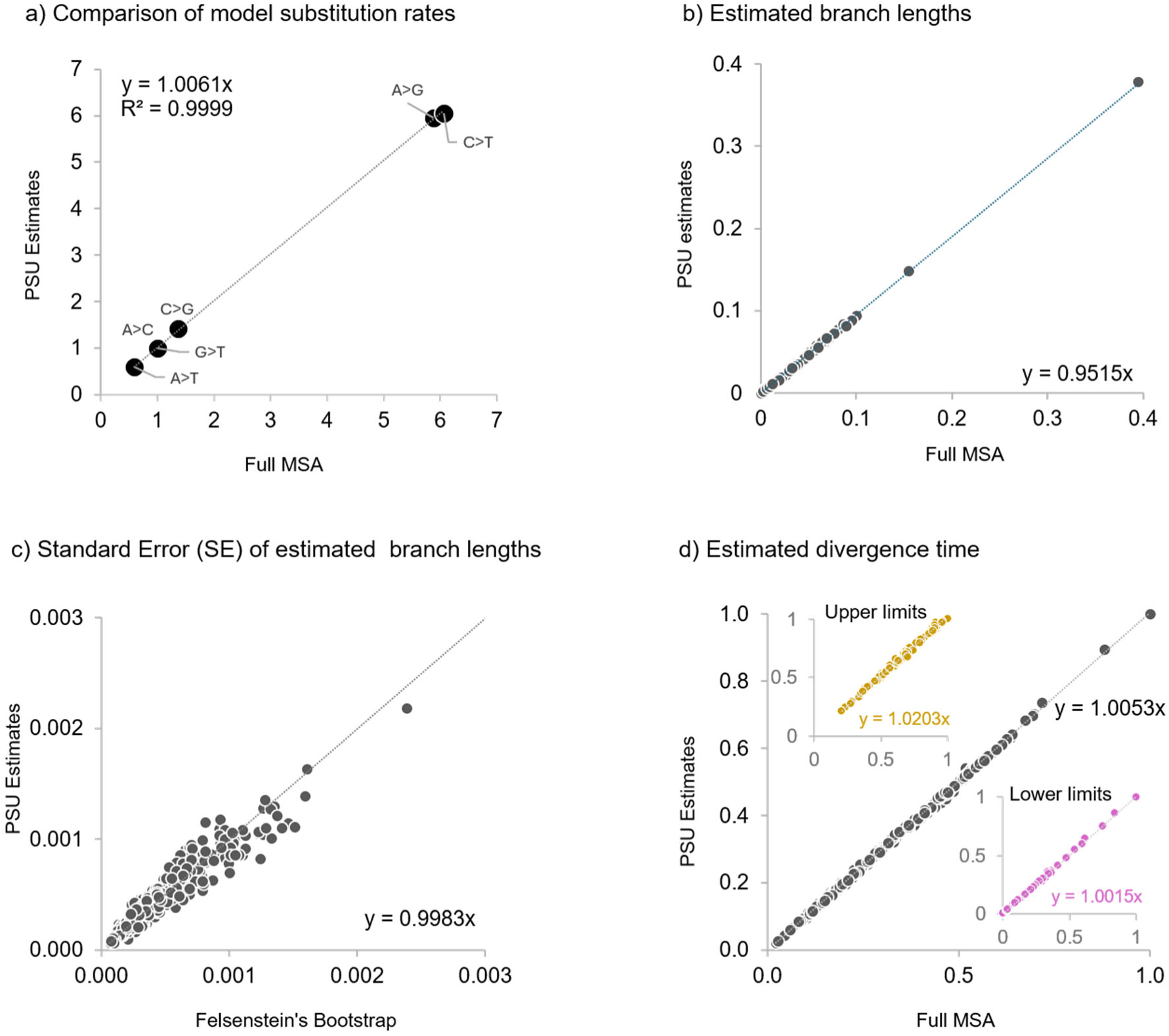
PSU estimates of model parameters, branch lengths, and divergence times for the 200×394 dataset. (**a**) Comparison of model substitution rate parameter estimates obtained from PSU and full dataset analyses. (**b**) Comparison of branch lengths for all internal and tip branches estimated using PSU and full dataset analysis. Branch lengths are estimated for the bird phylogeny and the GTR+G4 model of substitutions. (**c**) Comparison of branch length standard error (square root of estimated variance) from the standard bootstrap (x-axis) and PSU approach (y-axis). (**d**) Comparison of the relative time estimated using the RelTime approach for the given phylogeny with estimated branch lengths. The gray circle represents the relative node times, while the purple and yellow circles represent the upper and lower limits of 95% confidence intervals. All analyses were conducted using IQTREE 2 with and without PSU.

#### Estimating branch lengths and their variances

Branch lengths are central to molecular evolutionary analysis because they quantify evolutionary change along lineages and are used to identify rate variation, test evolutionary hypotheses, and estimate divergence times. In large phylogenomic alignments, estimating branch lengths by maximum likelihood can be computationally demanding because branch lengths and model parameters must be optimized across all site patterns. Their estimation using PSU provides a stringent test of its usefulness as a framework, as branch lengths are estimated directly from sequence variation rather than from resampling distributions.

A PSU protocol similar to that used for substitution-model parameters can be applied to estimation of branch lengths of a given phylogeny, where the initial *n* is determined using the empirical formula in Sharma and Kumar (2022). Here, the convergence of estimates can be assessed by monitoring the stability of short branch-length estimates as the subsample size increases (2*n*, 3*n*, …). Short branches provide a stringent target because they are harder to estimate reliably than long branches. In our analyses, a stopping rule based on a correlation greater than 0.99 for the shortest 25% of branch lengths between successive iterations was effective.

Applied to the 200×394 dataset, this protocol produced branch-length estimates that closely matched those from the full alignment (**Fig. 9b**; slope = 0.95, correlation = 0.99), while reducing runtime and memory usage by 4.6-fold and 10.8-fold, respectively. PSU can also estimate branch length variances, which are needed for hypothesis testing (Dopazo 1994) and for confidence intervals in divergence-time methods such as RelTime (Tamura et al. 2012; Tamura et al. 2018; Tao et al. 2020). After a stable subsample size is identified for branch length estimation, multiple subsamples and upsampling replicates are analyzed.

Branch-specific variances are calculated within each subsample and then averaged across subsamples, following Kleiner et al. (2014). Convergence is assessed using the coefficients of variation for the shortest 25% of branches; estimates are considered stable when the correlation between successive iterations exceeds 0.99. For the 200×394 dataset, this approach closely matched full-bootstrap variance estimates while requiring substantially less time and memory (**Fig. 9c**; **Table 1**).

#### Estimating divergence times and confidence intervals

Divergence time estimation represents one of the most computationally intensive applications in evolutionary biology because branch lengths, substitution rates, and calibration constraints must be considered simultaneously. Consequently, analyses of large phylogenomic alignments often require substantial computational resources and extended runtimes. The successful application of PSU to divergence time estimation would therefore demonstrate that the framework can accelerate not only phylogenetic reconstruction but also downstream evolutionary analyses that rely on phylogenomic trees.

Indeed, divergence times obtained using the RelTime framework (Tamura et al. 2012; Tamura et al. 2018; Tao et al. 2020) using branch lengths estimated from the PSU approach were highly concordant with those estimated from the full dataset (**Fig. 9d**; slope = 1.01, correlation = 0.99). In addition to the relative node time, their 95% confidence intervals from the PSU approach closely matched those obtained from the full-dataset analysis (**Fig. 9d**, CI_lower_: slope = 1.00 and correlation = 0.99; CI_upper_: slope = 1.02, and correlation = 0.99).

Overall, results from this section demonstrate that PSU is not limited to estimating confidence measures or selecting among competing models. The same computational strategy can also recover continuous evolutionary parameters and their associated uncertainties. This extension from support estimation to parameter estimation substantially broadens the scope of PSU and reinforces its role as a general framework for scalable phylogenomic inference.

### Bootstrap Consensus Phylogeny and Branch Lengths using PSU

The PSU framework can also support an integrated phylogenomic workflow, in which the upsampled replicate trees used to estimate clade support are also used to obtain a consensus phylogeny and produce branch lengths and substitution parameter estimates. The consensus of bootstrap replicate trees is often used as the inferred phylogeny (Felsenstein 1985). Because phylogenomic datasets frequently yield strong support for most clades, PSU-derived replicate trees can likewise be used to construct majority-rule consensus phylogenies. In such cases, clade hypotheses, support values, and estimates of phylogenetic heterogeneity are obtained in a single computational analysis.

Branch lengths and their variances can also be estimated directly from the PSU replicate trees used for building the consensus tree. For each branch in the consensus phylogeny, corresponding branch lengths are extracted from replicate trees for each subsample, and their means and variances are subsequently aggregated across subsamples. To avoid unstable estimates, variance calculations can be restricted to clades that occur in the fewest replicate trees within a subsample.

Tests using empirical datasets showed that branch-length estimates (slope = 0.95, correlation = 0.99) and branch-length variances (slope = 0.97, correlation = 0.98) closely matched those obtained from full-data analyses. These results highlight an important practical advantage of PSU: a single analysis can provide a consensus phylogeny, clade-support measures, branch-length estimates, model parameters, and associated uncertainty measures, thereby reducing the need for multiple separate full-data analyses in large phylogenomic studies.

### Applying PSU Beyond ML and Species Trees

The examples above have focused primarily on ML analyses of concatenated phylogenomic nucleotide sequence alignments using IQ-TREE 2 (see **Data and Code availability**). So, all the performance comparisons are direct, as they were conducted with and without PSU using IQ-TREE 2. The PSU framework can be implemented using other software with ML analysis engines, such as RaxML and MEGA. In fact, PSU implementation in MEGA is now underway (Kumar et al., in preparation).

Furthermore, PSU naturally applies to amino acid sequence alignments. For example, Sharma and Kumar (2022) analyzed multiple simulated and empirical protein sequence alignments. They reported that PSU analysis could select the same best-fit substitution model as the full MSA analysis using less than 1% of the site patterns for long phylogenomic alignments. For alignments containing only a few thousand site patterns, PSU is not expected to offer significant computational savings (Sharma and Kumar 2022). In fact, the initial value of *n* predicted using the empirically derived formula in Sharma and Kumar (2022) suggests not using PSU.

Sharma and Kumar (2025) applied PSU to estimate *NBS* for a long fungal amino acid sequence alignment. PSU required only ∼2% of the sites (11,210 sites from 609,772), and recovered all highly supported clades identified in the full MSA analysis. Interestingly, PSU also identified clades that received falsely high *FBS* values due to data errors or contentious phylogenetic relationships between concatenation-based and multispecies coalescent approaches, e.g., (Shen et al. 2017; Sharma and Kumar 2024; Sharma and Kumar 2025). Therefore, the PSU approach can also be used for amino acid sequence alignments, and the approach for predicting the initial number of patterns to subsample can be used to determine when its application is likely to be ineffective.

The PSU principle is also not inherently tied to the ML. Generally, PSU is useful whenever computational cost increases with alignment size while statistical power accumulates through the addition of sites and substitutions. Under these conditions, the PSU strategy can potentially reduce computational burden while preserving much of the information needed for inference. Maximum parsimony satisfies this requirement, as computational effort increases with alignment size and repeated site patterns need not be evaluated independently. Distance-based methods, including Neighbor-Joining (Saitou and Nei 1987), may also benefit when distance estimation or bootstrap testing must be repeated many times for long alignments. Bayesian phylogenetic analyses (Holder and Lewis 2003; Yang and Rannala 2012) are particularly interesting candidates for PSU because repeated likelihood evaluations during posterior sampling often dominate computational cost. Although the statistical behavior of PSU in Bayesian settings remains to be investigated, upsampled subsamples may provide useful approximations for estimating posterior support, branch lengths, and other evolutionary parameters.

The same logic may extend beyond species-level phylogenies. Large population-genomic datasets containing millions of polymorphic sites are increasingly used to estimate evolutionary relationships, demographic parameters, and measures of genetic diversity (Ellegren 2014; Luikart et al. 2018). Such datasets pose computational challenges similar to those encountered in species phylogenomics, suggesting that PSU-inspired approaches are likely to be useful, though modified adaptive parameter-tuning strategies may be required. Systematic evaluation of these applications remains an important direction for future research.

### Scaling with Increasing Numbers of Sequences

Until now, we have discussed how PSU addresses the computational burden in phylogenomics that arises from an increasingly larger number of sites. However, the number of sequences (taxa) is also growing rapidly, creating additional computational challenges (Pattengale et al. 2010; Minh et al. 2013; Sharma and Kumar 2021). Numerous methods have, therefore, been developed to accelerate phylogenetic inference when large numbers of taxa are analyzed, including rapid bootstrap procedures, improved tree-search algorithms, and approximate likelihood calculations (Stamatakis et al. 2008; Price et al. 2010a; Minh et al. 2013; Hoang et al. 2018). Importantly, these approaches address a different computational bottleneck than PSU, accelerating tree search and supporting estimation while still analyzing the full alignment. In contrast, PSU reduces the complexity of the alignment itself by limiting the number of unique site patterns evaluated via analysis of subsamples. Consequently, other approaches are complementary to PSU and can be combined with it to achieve computational gains in both dimensions of a phylogenomic dataset: many sites and many taxa.

For example, Sharma and Kumar (2021) estimated subsample-wise clade support using the Ultrafast Bootstrap (UFBoot) method (Minh et al. 2013; Hoang et al. 2018). Each subsample was analyzed using UFBoot, which repeatedly upsamples the subsample *R* times, typically *R* ≥ 1000, to generate replicate phylogenies from which clades and their corresponding subsample bootstrap support values, *bcl*, are estimated. These clade-specific *bcl* values are then used to estimate *BCL* and *NBS*. For a mammalian dataset containing 37 species and 1,391,742 sites, the combined PSU+UFBoot approach required only 0.83 hours and 0.16 GB of memory on a standard multi-core computer using 10 subsamples and default UFBoot settings (Sharma and Kumar 2021). This represented a substantial improvement over using UFBoot alone, which required 7.50 hours and 6.7 GB of memory, and PSU alone, which required 18.9 hours. Similar to the *FBS* results, all clades except one were recovered with *BCL* ≥ 95%, whereas the remaining clade received *BCL* = 90%. These results illustrate that combining PSU with fast bootstrap heuristics can yield multiplicative gains in runtime and memory efficiency.

Other advances in phylogenetic inference can likewise be integrated into the PSU framework, including improved tree-search strategies, approximate likelihood calculations, and alternative measures of branch support such as transfer bootstrap expectation (Lemoine et al. 2018). PSU can be paired with a wide range of inference engines provided that the computational burden is a direct function of the number of unique site patterns analyzed. PSU should therefore be viewed as complementary to ongoing algorithmic advances for large phylogenomic datasets rather than as an alternative to them.

### Limitations of the PSU framework

PSU is a theoretically motivated approximation rather than an exact reformulation of full-data phylogenetic inference. Its connection to the bag-of-little-bootstraps framework provides statistical motivation, but it does not establish equivalence between PSU and full-data analysis. At present, the strongest support for PSU comes from empirical performance across simulated and empirical phylogenomic datasets. PSU should therefore be viewed as a complementary framework that prioritizes scalability while approximating quantities that would otherwise require more expensive full-data analyses.

The effectiveness of PSU depends on whether relatively small subsamples capture representative patterns of evolutionary change to reconstruct all the evolutionary relationships. When datasets contain one or more taxa with extensive missing data, larger subsamples will be required to obtain stable bootstrap estimates for some clades. In such cases, the computational advantage of PSU will diminish, as a large fraction of sites will need to be included in the subsample to ensure that a sufficient number of substitutions are included for correctly reconstructing phylogenetic relationships of data-poor taxa. In such situations, one may harness PSU’s efficiency by, if possible, excluding data-poor taxa and then using phylogeny placement approaches to add them to the PSU phylogeny of the rest of the taxa.

PSU may also produce bootstrap support values that differ significantly from *FBS* when phylogenetic signals are concentrated in a few small subsets of highly informative sites. Because PSU relies on random site subsampling, some subsamples may omit these sites entirely, and *BCL* may underestimate *FBS* with small subsamples, requiring large subsamples to reproduce *FBS*. At the same time, the discrepancy may indicate that support for a clade is driven by overly influential, likely disruptive sites (Sharma and Kumar 2025). In this sense, PSU can be useful as a diagnostic framework for situations in which information from a few sites drives phylogenetic signal in the analysis of concatenated alignments, e.g., (Brown and Thomson 2016; Shen et al. 2017; Walker et al. 2018; Sharma and Kumar 2024).

Because PSU relies on random site subsampling, results can vary across independent runs, particularly when the number of subsamples is small or when subsample sizes are too small to capture representative evolutionary signals. This variability can affect estimates of clade support and other evolutionary inferences. Such run-to-run variation is common in other stochastic phylogenetic procedures, including bootstrap analyses, Bayesian posterior sampling, and heuristic tree searches. e.g., (Shen et al. 2020; Kumar et al. 2023). In PSU, such variability can be reduced by increasing the number of subsamples, increasing subsample size, or both. Adaptive protocols also attempt to mitigate this problem by expanding the analysis until estimates stabilize, although optimal settings remain dataset-dependent.

Like other phylogenetic methods, PSU inherits sensitivities associated with the underlying inference procedure applied to a concatenated sequence alignment. Model misspecification, compositional heterogeneity, long-branch attraction, or violations of other assumptions can affect PSU analyses in much the same way that they affect corresponding full-data analyses. PSU does not eliminate these challenges, although the distribution of estimates across subsamples may provide additional insight into the stability of the clades using concatenated datasets.

Finally, PSU does not replace methods that explicitly model particular biological processes. For example, *NBS* can reveal conflict in concatenated alignments, but it is not a substitute for multispecies coalescent approaches when the goal is to model gene-tree variation directly. Similarly, PSU is meant for long concatenated alignments, whose systematic evaluation of classes of evolutionary inference problems and methods is still needed to refine convergence criteria, identify failure modes, quantify computational trade-offs, and establish best practices for PSU analyses.

### Conclusions

PSU is a general strategy for scalable phylogenomic inference that analyzes many small representations (site subsamples) of a large concatenated alignment, upsamples them before inference, and aggregates the resulting estimates. Its effectiveness arises from a useful separation between computational burden and inferential power: computational cost is strongly influenced by the number of unique site patterns in the dataset, whereas statistical power and many measures of inferential precision depend primarily on the total amount of evolutionary information represented by sites and substitutions. By reducing the former while restoring the latter through upsampling, PSU can approximate a wide range of full-data analyses at substantially lower computational cost.

The PSU framework extends well beyond accelerating bootstrap support estimation. Because PSU produces distributions of support values across independent subsamples, it also provides direct information about concordant and conflicting phylogenetic signals that is often obscured when a single full-alignment value summarizes support. This makes PSU not only a computational shortcut but also a diagnostic framework for detecting heterogeneity in phylogeny signals across the dataset. Also, as shown above, the PSU strategy can be applied to substitution model selection, likelihood-based hypothesis testing, branch-length estimation, divergence-time analysis, and parameter variance estimation. This common computational principle unifies all current PSU applications despite their differing inferential objectives.

We have shown that a single PSU analysis can yield the consensus bootstrap phylogeny and support values, model parameters, branch lengths, and associated uncertainty measures, thereby avoiding separate full-alignment analyses. Adaptive procedures for selecting the subsample size, the number of subsamples, and the number of upsampling replicates further enhance the framework’s practicality and reproducibility. Furthermore, methods to speed up calculations for large numbers of sequences can be used alongside PSU to improve computational efficiency for large datasets, as discussed above.

Although this article focused primarily on maximum-likelihood phylogenomics of species, the PSU principle is not inherently tied to a particular inference method or taxonomic scope. We anticipate that the same principle will be useful for Bayesian, maximum parsimony, and distance-based phylogenetics, as well as for large-scale population polymorphism analyses, because the computational cost increases faster than the statistical information gained from additional sites. These opportunities suggest that PSU can serve as a broadly useful computational paradigm for evolutionary analysis of massive datasets.

As phylogenomic datasets continue to grow, scalable approaches will be essential for maintaining statistical rigor and reproducibility in evolutionary inference, while also ensuring accessibility on standard computing platforms. With upcoming implementation in a widely used software package, e.g., MEGA (Kumar et al. 2024), and growing interest in scalable phylogenomic methods, PSU can become an important component of future big-data evolutionary analyses.

## Data and Code availability

The phylogenomic datasets discussed in this article are publicly available from the corresponding references. R codes for performing PSU analyses are available on GitHub at https://github.com/ssharma2712.

## Acknowledgments

We thank John Allard and Glen Stecher for many helpful comments. This work was supported by a research grant from the National Institutes of Health (R35GM139540-06) and the National Science Foundation (DBI-2505985).

## Appendix

### Median versus Mean for Binary Scoring in Bootstrap Analysis

The little bootstrap framework requires aggregation of subsample-wise estimates. While Kleiner et al. (2014) focused primarily on mean bagging, Sharma and Kumar (2021) found that median bagging yields estimates closer to conventional bootstrap support. To illustrate why median bagging is appropriate, we considered an indicator-bootstrap framework analogous to a phylogenetic bootstrap analysis. In this setting, each bootstrap replicate receives a binary score: 1 if it supports a hypothesis and 0 otherwise. The bootstrap estimate is then the proportion of replicates that have a score of 1. This parallels phylogenetic bootstrapping, where a clade receives a score of 1 when present in a replicate tree and 0 when absent.

We generated data from a normal distribution and compared p-values obtained using the standard bootstrap with those obtained using the bag-of-little-bootstraps framework. For the little bootstrap analyses, subsamples of size (*N^g^*) were analyzed using both mean and median bagging. Median bagging consistently produced estimates that more closely matched those obtained from the standard bootstrap (**Fig. A1**). Mean bagging was more sensitive to a small number of subsamples that strongly supported or rejected the hypothesis. These extreme subsamples tend to pull the average toward 0.5, the expected value under random support. Consequently, mean bagging tends to overestimate small *P*-values and underestimate large p-values. A similar pattern was observed for phylogenetic bootstrap support values (Sharma and Kumar 2021), where median bagging yielded *BCL* values that more closely approximated conventional Full-data bootstrap support (*FBS*).

**Figure A1.**
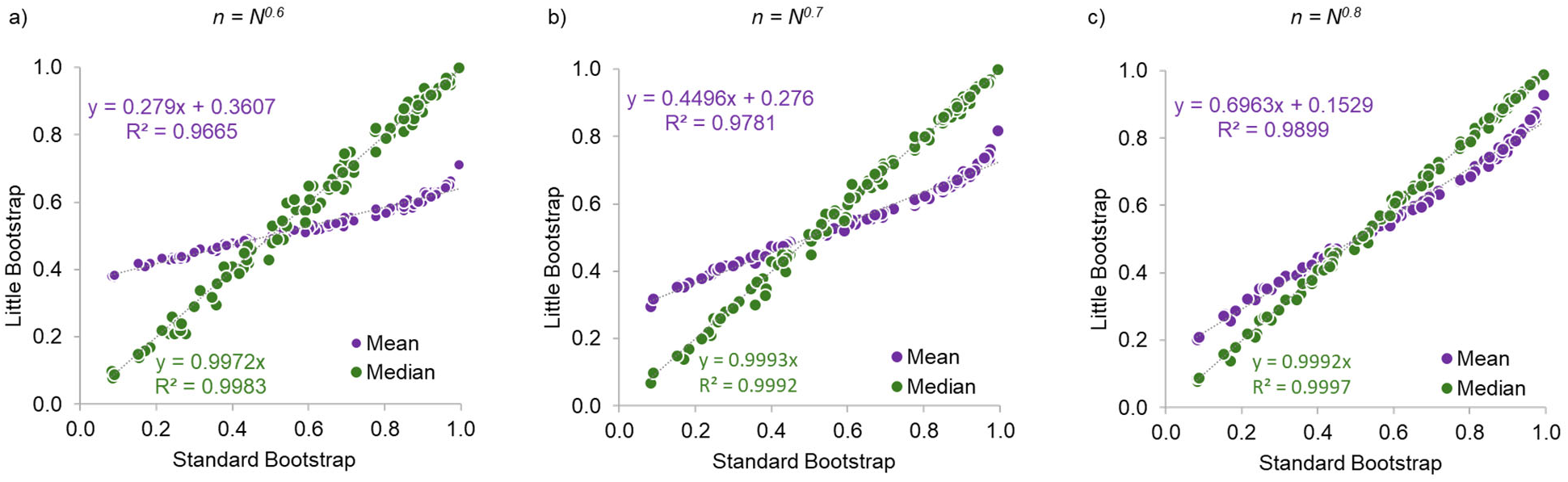
Comparison of p-values estimated by the standard and little bootstrap with mean (blue) and median bagging (orange). Data were simulated from a normal distribution X ∼ *Normal*(μ = −5, σ = 1). We generated 100 random samples of size *N* = 25,000 and tested H_0_: μ = −5.01 against the one-sided alternative H_1_: μ < −5.01, a threshold chosen to produce a broad range of p-values. For each sample, p-values were estimated using the standard bootstrap and the bag-of-little bootstraps. In the little bootstrap, subsamples of size *n* = *N^g^* were drawn, with *g* = 0.6, 0.7, or 0.8 in panels **a**-**c**, respectively. For each value of *g*, 1000 subsamples were analyzed, and 100 upsampled replicates were generated from each subsample. Subsample-wise p-values were then aggregated using either the mean or the median.

**Supplementary Table S1.** Information on datasets shown in. **Figure 1**.

| Data | Base type | Taxa | Total Sites | Unique Site patterns (*U*) | Data Size (*D*) | Median pairwise difference (%) (Minimum - Maximum) | DOI |
| --- | --- | --- | --- | --- | --- | --- | --- |
| Insects A | DNA | 174 | 3,011,544 | 2,045,783 | 355,966,242 | 13.9 (0.3 - 30.1) | https://doi.org/10.1016/j.cub.2017.01.027 |
| Butterflies | DNA | 61 | 5,267,461 | 3,762,723 | 229,526,103 | 21.9 (1.1 - 28.3) | https://doi.org/10.1093/sysbio/syz030 |
| Vertebrate | AA | 58 | 1,806,035 | 1,547,914 | 8,977,9012 | 22.4 (2.1 - 30.3) | https://doi.org/10.1093/sysbio/syv059 |
| Plants A | AA | 360 | 19,449 | 190,352 | 68,526,720 | 3.8 (0.0 - 18.9) | https://doi.org/10.1186/1471-2148-14-23 |
| Insects B | DNA | 48 | 2,938,039 | 1,396,402 | 67,027,296 | 15.1 (3.2 - 21.1) | https://doi.org/10.1016/j.ympev.2017.12.005 |
| Fungi | AA | 86 | 609,772 | 562,771 | 48,398,306 | 51.2 (0.4 - 59.7) | https://doi.org/10.1038/s41559-017-0126 |
| Birds | DNA | 200 | 394,684 | 226,490 | 45,298,000 | 7.4 (0.5 - 16.6) | https://doi.org/10.1038/nature15697 |
| Mammal A | DNA | 37 | 1,391,742 | 775,579 | 28,696,423 | 12.8 (0.5 - 26.5) | https://doi.org/10.1073/pnas.1211733109 |
| Yeast | AA | 23 | 634,530 | 390,960 | 8,992,080 | 41.9 (5.7 - 52.6) | https://doi.org/10.1038/nature12130 |
| Rodents | DNA | 37 | 1,207,638 | 96,293 | 3,562,841 | 2.4 (0.2 - 5.8) | https://doi.org/10.1093/sysbio/syz044 |
| Mammal B | DNA | 274 | 7,370 | 4,303 | 1,179,022 | 15.2 (0.3 - 22.2) | https://doi.org/10.1098/rspb.2012.0683 |
| Plants B | DNA | 16 | 4,246,454 | 17,789 | 284,624 | 4.8 (0.4 - 15.4) | https://doi.org/10.1016/j.ympev.2018.08.011 |
| Lassa Virus | DNA | 179 | 3,186 | 1,475 | 264,025 | 21.1 (0.0 - 24.7) | https://doi.org/10.1016/j.cell.2015.07.020 |

**Note.** Data refers to genomic sequence alignments from the indicated species. Bases denote the type of nucleotide or amino acid data. Dataset size (*D*) was calculated as the product of the number of taxa and the number of unique site patterns (*U*), because both factors are associated with the computational time and memory required for maximum-likelihood analysis. The median, minimum, and maximum pairwise sequence difference per site represent the level of sequence diversity within each alignment. DOI provides the digital object identifiers of the studies in which these datasets were originally published and analyzed.

